# A conserved oomycete CRN effector targets and modulates tomato TCP14-2 to enhance virulence

**DOI:** 10.1101/001248

**Authors:** Remco Stam, Graham B. Motion, Petra C. Boevink, Edgar Huitema

**Author notes:** Correspondence: Edgar Huitema.

## Abstract

*Phytophthora* spp. secrete vast arrays of effector molecules upon infection. A main class of intracellular effectors are the CRNs. They are translocated into the host cell and specifically localise to the nucleus where they are thought to perturb many different cellular processes. Although CRN proteins have been implicated as effectors, direct evidence of CRN mediated perturbation of host processes has been lacking. Here we show that a conserved CRN effector from *P. capsici* directly binds to tomato transcription factor SlTCP14-2. Previous studies in *Arabidopsis thaliana* have revealed that transcription factor TCP14 may be key immune signalling protein, targeted by effectors from divergent species. We extend on our understanding of TCP targeting by pathogen effectors by showing that the *P. capsici* effector CRN12_997 binds to SlTCP14-2 in plants. SlTCP14-2 over-expression enhances immunity to *P. capsici*, a phenotypic outcome that can be abolished by co-expression of CRN12_997.

We show that in the presence of CRN12_997, SlTCP14-2 association with nuclear chromatin is diminished, resulting in altered SlTCP14 subnuclear localisation. These results suggest that CRN12_997 prevents SlTCP14 from positively regulating defence against *P. capsici*. Our work demonstrates a direct interaction between an oomycete CRN and a host target required for suppression of immunity. Collectively, our results hint at a virulence strategy that is conserved within the oomycetes and may allow engineering of resistance to a wide range of crop pathogens.

## Introduction

Plant-microbe associations feature dynamic interplay and engagement of vast signalling networks that help determine interaction outcomes. Besides preformed barriers, plants deploy surface exposed Pattern Recognition Receptors (PRRs) to perceive Pathogen or Microbe Associated Molecular Patterns (P/MAMPs) and mount an immune response. In the vast majority of cases, PAMP perception leads to activation of complex signalling cascades, cellular defences and transcriptional reprogramming events, that lead to Pattern Triggered Immunity (PTI) and resistance to microbes (1). Considering that plants are exposed to dynamic microbial communities, have to fend off potential pathogens whilst facilitating beneficial associations with microbes throughout their growth cycle; plants require a finely tuned host immune signalling network. It is therefore not surprising that recent efforts to map the Arabidopsis immune signalling network, has resulted in the identification of a vast and complex set of interconnected cascades that help explain how appropriate immune responses are mounted against diverse microbes (2, 3).

Pathogen ingress implies that the host immune signalling network is perturbed and as a consequence, immunity is compromised. Pathogens thus are specialised microbes that have acquired and evolved the ability to suppress immune responses in their respective host. Indeed, genome sequencing, functional genomics and detailed biochemical analyses have helped identify secreted proteins (effectors) in most if not all pathogens. Effectors that are secreted during infection either accumulate and function in the host apoplast (apoplastic effectors) or translocate inside host cells where they travel to distinct subcellular compartments and target resident host proteins (cytoplasmic effectors). Modification of cellular processes associated with immunity are widely regarded as the principal mechanism by which effectors ‘trigger’ susceptibility to pathogens (ETS, Effector-Triggered Susceptibility (4, 5)) although a mechanistic understanding of target perturbation is lacking in most pathosystems. To understand pathogen virulence strategies, or more specifically, the role(s) of effectors towards parasitism, the identification of their host targets represents a critical step. Consequently, recent efforts have focused on target identification and subsequent studies on the host processes affected, which in some cases have unveiled molecular events underpinning immunity and susceptibility with unprecedented new detail.

Systems levels analyses of effector-target interactions and immune signaling network topology have provided additional insights into how diverse pathogens can target key immune regulators (hubs) to achieve virulence. A matrix two-hybrid screen using effector proteins from oomycete *Hyaloperonospora arabidopsidis* and *Pseudomonas syringae,* against a large protein set from *Arabidopsis thaliana* demonstrated that effector arsenals encoded by diverse pathogens, share common host targets. Furthermore, common host targets often formed important hubs in the *Arabidopsis* immune network, suggesting that pathogens shaped their effector repertoires to target important regulators and suppress immunity during infection (6). One such example is the Arabidopsis transcription factor TCP14. *At*TCP14 was identified as a major hub targeted by *H. arabidopsidis* and *Ps. syringae* effectors (6, 7). TCP proteins form a large family of transcription factors with a DNA binding basic helix loop helix domain, dubbed the TCP domain (8). Although *At*TCP14 is thought to play various roles in plant growth and development (9–12), TCP-family members have been implicated as regulators of plant immunity. Phytoplasma Aster Yellows, strain Witches’ Broom (AY-WB), effector SAP11 destabalises CINCINNATA (CIN)-related TCPs which leads to a compromised JA response and enhanced fecundity of its insect vector *Macrosteles quadrilineatus*, facilitating the transmission and spread of AY-WB (13). Chandran *et al.* (14) suggest that TCPs might play a role during infection of Powdery Mildew *Golovinomyces orontii.* These observations suggest that divergent pathogens have independently evolved the ability to target important host proteins to suppress immunity. If true, conservation of important effector-target interactions may form another important means besides convergent evolution, by which targeting of important host proteins by divergent pathogen effectors, is achieved.

Filamentous plant pathogens, including oomycetes, are ranked amongst the greatest threats to plants, animals and ecosystems (15). Amongst the oomycetes, *Phytophthora infestans* and *P. sojae* form major threats to potato and soybean production respectively (16, 17). *P. capsici* has emerged as a significant threat to dozens of valuable crops, including cucumber, pepper and tomato (18, 19). *Phytophthora* spp are hemi-biotrophic pathogens. Infection features an early biotrophic stage, during which host tissue appears healthy, followed by a necrotrophic phase marked by cell death and tissue collapse. Infection is coupled to two massive transcriptional changes in both host and pathogen, collectively thought to underpin disease development and pathogen lifestyles on hosts (8).

*Phytophthora* genomes are thought to encode at least two major classes of intracellular effectors that are thought to contribute to virulence. RXLR effectors are named after their N-terminal Arg-x-Leu-Arg motif, whereas CRN (For <u>Cr</u>inkling and <u>N</u>ecrosis) proteins all carry an LFLAK-motif (Leu-Phe-Leu-Ala-Lys) in their N-terminus (20, 21). Members of both effector families are modular proteins with conserved N-termini thought to specify translocation whereas C-terminal domains carry effector activities. Intriguingly, CRNs proteins are more widespread in the oomycetes than the RXLR effectors class. CRN proteins have been found in all plant pathogenic oomycetes sequenced to date, whereas RXLR coding genes only appear to be present in members of the *Peronosporales* lineage (22–27). In addition, CRN N-terminal regions, along with some of their C-terminal counterparts, are highly conserved amongst plant pathogenic oomycetes, suggesting basic roles in oomycete biology or parasitism (28). Although CRN effectors are commonly found in plant pathogenic oomycetes, little is known about their contribution towards virulence or their activities. Localisation of CRN C-termini in plants revealed that a diverse array of effector domains from a wide range of oomycetes, accumulate in nuclei. Some CRN domains cause cell death and chlorosis phenotypes in plants, suggesting perturbation of cellular processes by the effector activities carried by CRN C-termini (28–31). More detailed analyses revealed that in the case of the *P. infestans* CRN8 effector domain, its accumulation in the nucleus is required for cell death. Additionally, PiCRN8, has been shown to possess kinase activity *in planta*, firmly confirming the presence of molecular functions that could contribute to the infection process *in vivo* (32). Despite these advances, critical information about the processes that are targeted by this protein family is missing.

Here we show that *Phytophthora capsici* CRN12_997 interacts with the tomato transcription factor SlTCP14-2 *in planta*. Over-expression of SlTCP14-2 enhances immunity to *P. capsici*, a phenotype that can be reversed upon co-expression of CRN12_997 but not by the related effector CRN125_11. Consistent with a function towards altering SlTCP14-2 function, we present evidence suggesting that CRN12_997 binds a complex containing SlTCP14-2 and causes its re-localisation in the nucleoplasm and nucleolus. Importantly, we show that the presence of CRN12_997 lowers the levels of SlTCP14-2 bound to chromatin, possibly decreasing SlTCP14 stability and underpinning re-localisation. By demonstrating that a specific effector-target interaction is present in two divergent oomycete-host interactions, we suggest that conservation of ancient host-target interactions is an additional mechanism by which common virulence strategies in pathogens emerge.

## RESULTS

### Conserved *P. capsici* CRN effectors may target transcription factor TCP14

We reported that the CRN effector repertoire is widespread amongst plant pathogenic oomycetes and that a subset of these CRN effector domains is present in most sequenced genomes (28). The DXX C-terminal domain is among the most conserved domains and in *Hyaloperonospora arabidopsidis* three DXX domain-containing effectors have been identified. A matrix Yeast two Hybrid screen against a large subset of the Arabidopsis proteome identified the transcription factor TCP14 as a candidate target for all three effectors (6). AtTCP14 is targeted by effectors from a growing number of diverse pathogens suggesting that this transcription factor is a key regulator of the Arabidopsis immune network. Given that the *P. capsici* genome encodes DXX containing CRN effectors and AtTCP14 may have orthologous genes in the tomato genome, we asked whether CRN-TCP14 interactions occur within the *P. capsici*-tomato patho-system.

We identified two effectors featuring a DHB-DXX-DHA domain configuration amongst our set of 84 *P. capsici* CRN candidate genes (Figure S1A). Comparisons between both PcCRN12_997 and PcCRN125_11 and their possible *H. arabidopsidis* counterparts (HaRxLCRN4, 14 and 17), revealed 41% sequence identity in pairwise alignments between DXX domains. To assess relationships between CRN DXX domains in different species, we made a phylogenetic reconstruction of the C-terminal domains. This shows that CRN coding genes group together in species-specific clusters (Figure S1B), suggesting that these genes arose before speciation. If CRN-target interactions are both ancestral and preserved, this could suggest that *P. capsici* CRN12_997 or CRN125_11 binds AtTCP14 orthologs during infection of its hosts.

Blast analyses identified two putative tomato orthologs of AtTCP14, which we named SlTCP14-1 and SlTCP14-2. Both share high sequence similarity with AtTCP14, though both proteins are more similar to each other than to AtTCP14 (Figure S2). Both tomato TCP proteins contain the highly conserved Helix-loop-Helix motif, including the conserved Cysteine at the start of the first helix (33)(Figure S2A). To learn more about SlTCP14 gene expression, we examined microarray data from a *P. capsici* infection time course on tomato (34). These data show that SlTCP14-1 and SlTCP14-2 are expressed in tomato but down-regulated in the early infection time points (Figure S2B). These changes of gene expression coincide with the expression of DXX-domain containing CRNs as reported previously (28, 34). These results suggest that SlTCP14-1 and SlTCP14-2 may form important host targets whose expression is repressed during the interaction with *P. capsici*.

### BiFC shows YFP reconstitution with CRN12_997 and SlTCP14-2

Given the presence of orthologous sequences for both *H. arabidopsidis* effectors and their putative targets in *P. capsici* and tomato respectively, we aimed to test whether CRN12_997, CRN125_11 and both SlTCP14 proteins can interact *in vivo*. Bimolecular Fluorescence Complementation (BiFC) experiments, in which we co-expressed candidates fused to the N-terminal half of YFP (pCL112::CRN) and YFP C-terminus (pCL113::TCP) led to the specific accumulation of YFP signal in nucleoli for CRN12_997 and SlTCP14-2. We found low levels of YFP signal in the nucleoplasm for other CRN-TCP combinations (Figure 1). Given that we found YFP signal in multiple treatments, we quantified the level of specificity between CRN-TCP combinations. This revealed that in addition to specific localisation in the nucleolus, CRN12_997 and SlTCP14-2 co-expression led to a higher number of fluorescent nuclei (with YFP signal in the nucleolus) when compared to other combinations (Figure 1E). Although we found differences in protein abundance levels between the two CRN fusions (Supplemental Figure 3A), these do not correlate with the differences in YFP signal between BiFC treatments. Taken together these observations show that there is strong reconstitution of YFP fluorescence specifically with CRN12_997 and SlTCP14-2 treatments, suggestive of an interaction between these proteins *in vivo*.

**Figure 1.**
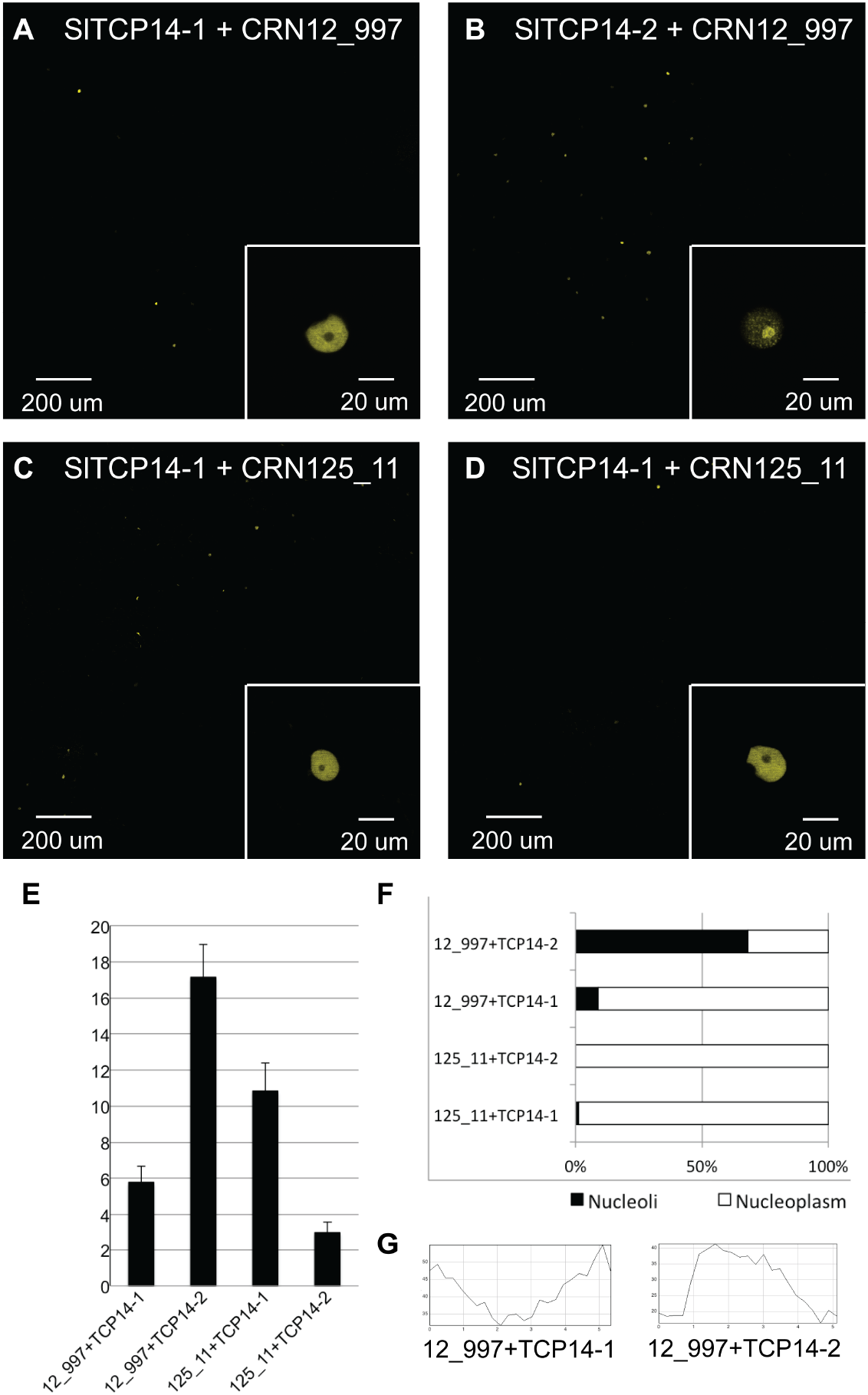
Split YFP confirms interaction of SlTCP14-2 and CRN12_997. A-D) Reconstituted YFP fluorescence is observed in all CRN/TCP combinations, however the number of fluorescent nuclei is significantly higher for CRN12_997 + SlTCP14-2 and with this combination reconstitution of fluorescence is observed in the nucleolus (insets). E) Amount of fluorescent nuclei per single slice image. F) Percentage of nuclei with fluorescence observed in the nucleolus. G) Fluorescence intensity profiles for insets of image A and B resp. showing relative higher fluorescence in the nucleolus of inset B.

### CRN12_997 and SlTCP14-2 specifically interact *in vivo*

To confirm our BiFC results we performed co-immunoprecipiation experiments using FLAG-tagged SlTCP14-2 and strepII-tagged EGFP-CRN fusion proteins. SlTCP14-2 was co-expressed with CRN12_997 and CRN125_11 fusions in *N. benthamiana* after which complexes were purified from total leaf extracts by using Mag-strep and FLAG M2 magnetic beads. Immunoprecipitation of CRN12_997, CRN125_11 fusion proteins and free (untagged) EGFP in the presence of SlTCP14-2, using strepII specific magnetic beads, led to a specific TCP signal for CRN12_997 only, suggesting a specific interaction between CRN12-997 and SlTCP14-2 proteins (Figure 2). Conversely, we were able to co-purify CRN12_997 in a FLAG pull down of SlTCP14-2, further confirming this interaction (Figure S4). From this, we conclude that consistent with our BiFC results, CRN12_997 specifically binds SlTCP14-2.

**Figure 2.**
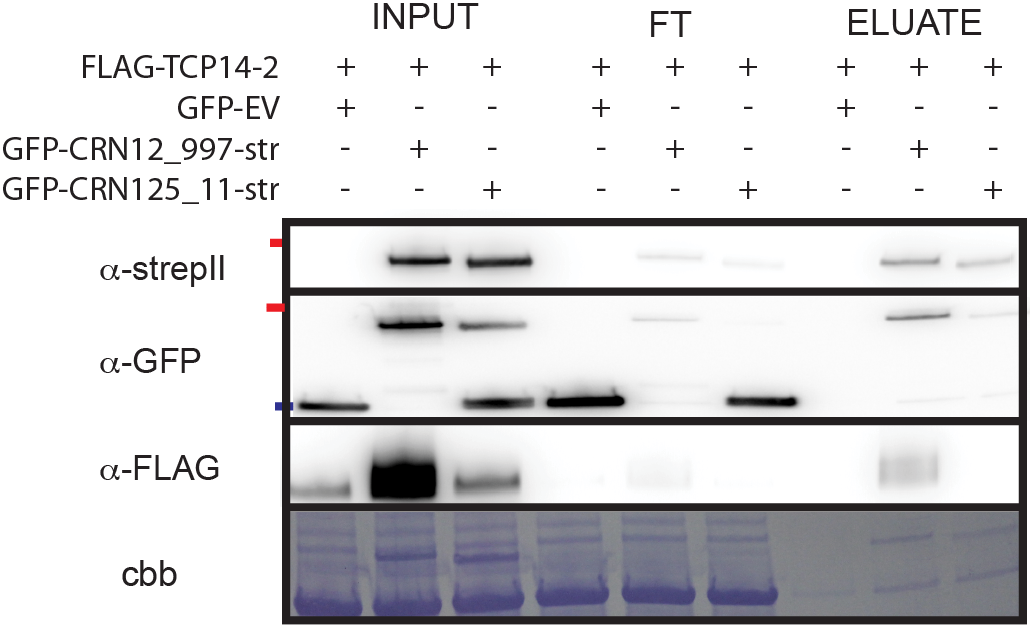
Co-immunoprecipitation shows a specific interaction between CRN12_997 and SlTCP14-2. StrepII-and GFP-tagged CRN proteins are present in all input samples and also visible after IP. FLAG-SlTCP14-2 is visible in the input samples and can only be observed in the eluate when CRN12_997 is co-expressed.

### SlTCP14-2 contributes to immunity to *P. capsici*

We asked whether SlTCP14-2 contributes to immunity as was suggested in for its *Arabidopsis* ortholog (6). We overexpressed SlTCP14-2 in *N. benthamiana* and infected infiltrated leaf panels with *P. capsici* zoospores as described previously (28). *P. capsici* infection of *N. benthamiana* leaves ectopically expressing SlTCP14-2 was found to be impaired as lesion growth rates were reduced when compared to empty vector controls (Figure 3A). This phenotype was specific to SlTCP14-2 expression since SlTCP14-1 expression did not significantly affect *P. capsici* growth when compared to the empty vector control (EGFP) (Figure 3A). Furthermore, expression of SlTCP14-2 inhibited *P. capsici* sporulation as sporangia were found to emerge only outside SlTCP14-2 expressing sites or in control sites (Figure 3C).

**Figure 3.**
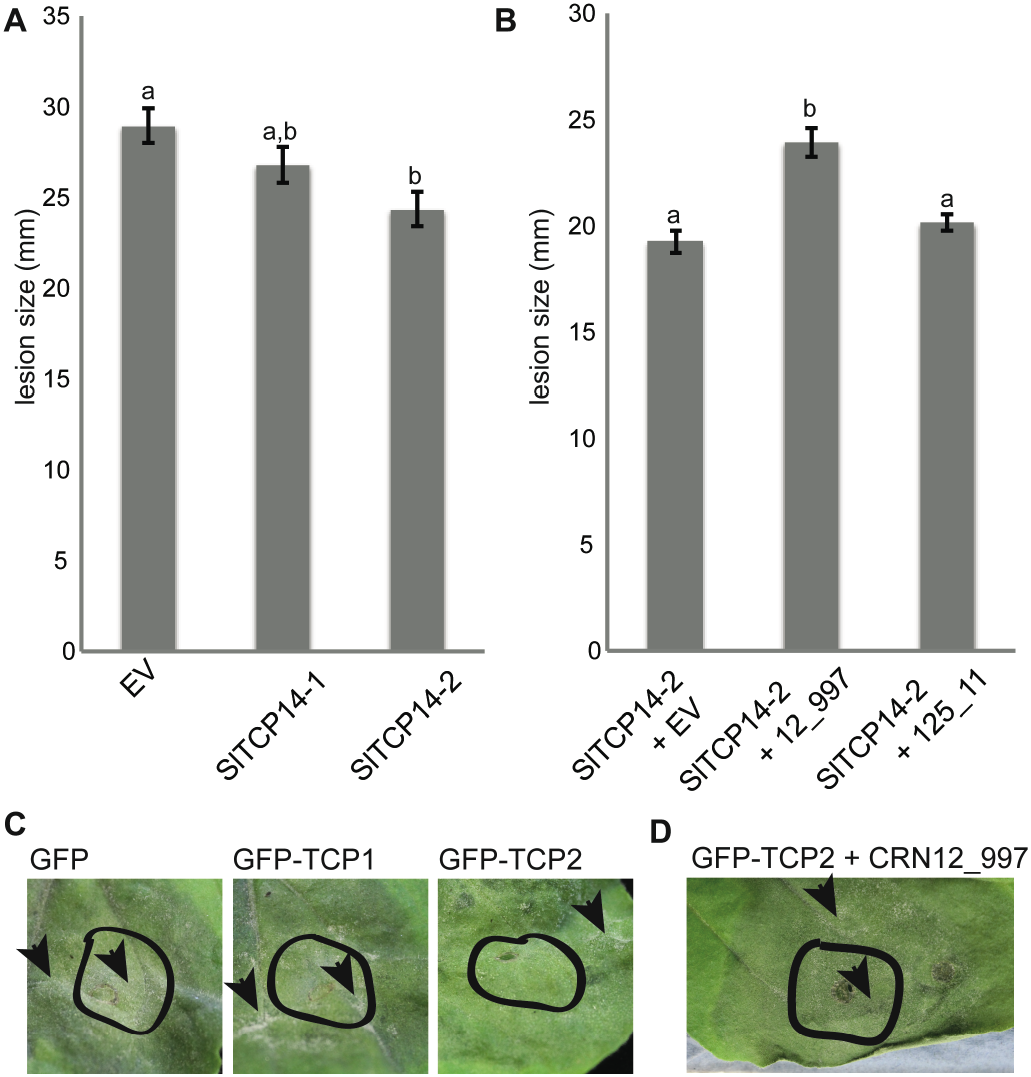
Over-expression of Sl-TCP14-2 reduces *P. capsici* virulence, co-expression of CRN12_997 is able to counteract. A) Over-expression of SlTCP14-2 in *N. benthamiana* shows a significant reduction in growth compared to over expression of free GFP (Anova p < 0.01) SlTCP14-1 shows an intermediate growth reduction. B) Co-expression of SlTCP14-2 with CRN12_997, but not with CRN125_11 partially restores growth retention. C) Expression of SlTCP14-2 delays spore formation, no sporangia (arrowheads) can be observed within the infiltrated site (black line) three days post infection. D) After co-expression of CRN12_997 with SlTCP14-2 sporangia can be observed at 3 dpi

### CRN12_997 counteracts SlTCP14-2 induced growth inhibition

Given that SlTCP14-2 over expression enhances immunity to *P. capsici*, we hypothesised that endogenous CRN12_997 is normally sufficient to suppress immunity, but high levels of SlTCP14-2 renders endogenous CRN12_997 levels inadequate. If true, ectopic expression of CRN12_997 should boost *P. capsici* growth in the presence of high SlTCP14-2 levels. We co-infiltrated SlTCP14-2 with CRN12_997, CRN125_11 and empty vector combinations, followed by *P. capsici* infection assays. These experiments showed that CRN12_997 over-expression reconstitutes growth to levels seen in empty vector controls. Significant growth enhancement (ANOVA, p < 0.01) was specific to co-expression of CRN12_997 with SlTCP14-2. CRN125_11 co-expression was found not to enhance growth in these assays when compared to the control (Figure 3B). As expected, we found that in the presence of CRN12_997, spore formation is also reconstituted despite over expression of SlTCP14-2 at three days after infection (Figure 3D). To show that these effects are due to the presence of fusion proteins rather then large differences in levels or stability, all constructs were expressed in *N. benthamiana* on multiple occasions and found to be stable when expressed individually (Figure 2 and S3).

### CRN12_997 alters subnuclear localisation of SlTCP14-2

Given that CRN12_997 and SlTCP14-2 localise and interact in the nucleus, we asked whether on the subnuclear level, CRN12_997 and SlTCP14-2 co-localise. We co-expressed CRN12_997 and SlTCP14-2, fused to EGFP and tagRFP respectively, in *N. benthamiana* and assessed localisation. Co-expression together with EGFP resulted in a speckled sub-nuclear distribution pattern for SlTCP14-2 two days after agroinfiltration (Figure 4A-C). Interestingly, co-expression of CRN12_997 with SlTCP14-2 abolished the characteristic speckled subnuclear localisation pattern (Figure 4D-F). Since co-expression of CRN125_11 did not affect SlTCP14-2 localisation, our results suggest that CRN12_997 binding of SlTCP14-2 alters localisation (Figure 4G-I).

**Figure 4.**
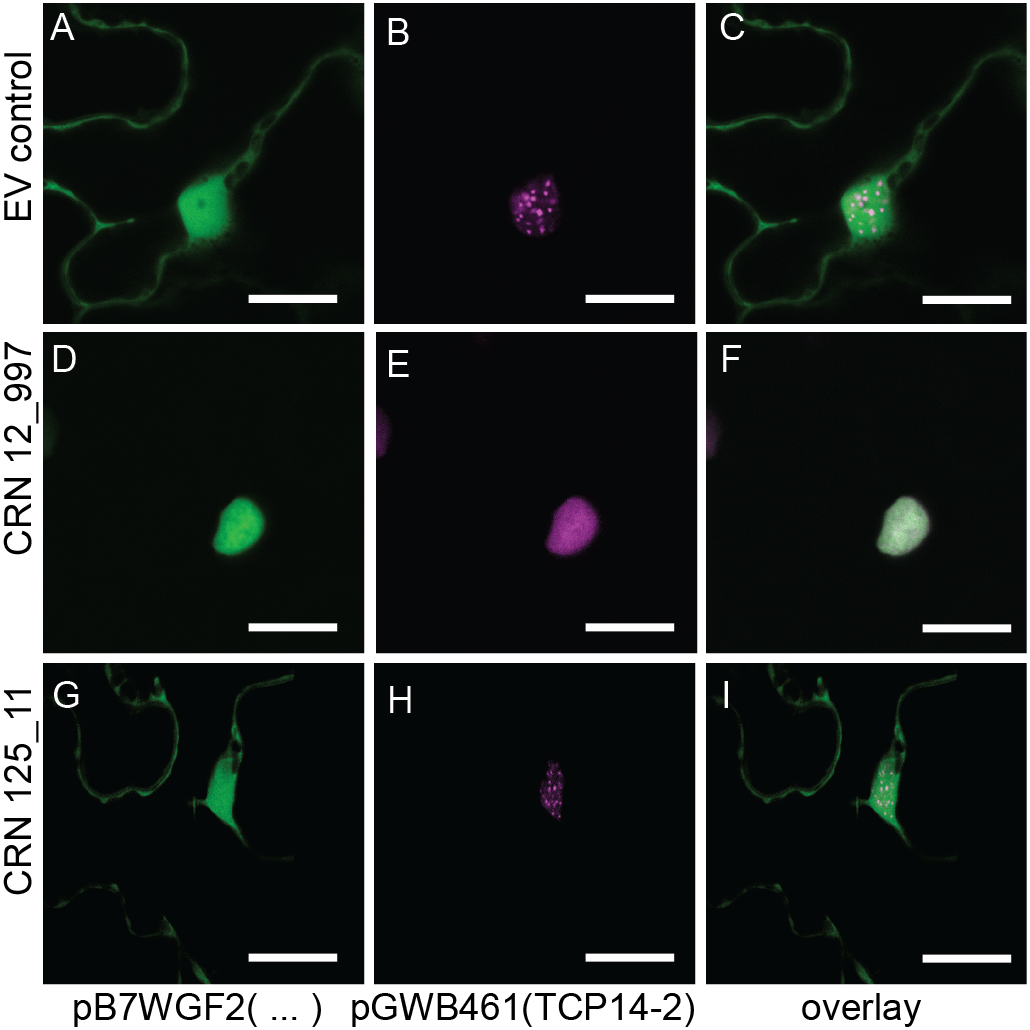
Co-expression of CRN12_997 and SlTCP14-2 results in altered TCP localisation. Over expressed RFP-tagged SlTCP14-2 in *N. benthamiana* plants shows a specific speckled phenotype when co-expressed with an eGFP control (A-C) or non-binding GFP-tagged effector CRN125_11 (G-I). Co-expression with GFP tagged CRN12_997 abolishes this pattern. Panels show GFP channel (A,D,G), RFP channel, (B,E,H) or overlay images (C,E,I). Scale bar is 10 um

To test whether relocalisation of SlTCP14-2 is a biologically relevant phenomenon, we assessed the localisation of transiently overexpressed SlTCP14-2 in transgenic *N. benthamiana* plants expressing an EGFP ER marker. Consistent with our co-localisation results, SlTCP14-2 shows diffuse nuclear localisation in cells that are infected with *P. capsici* (Figure 5). Assessment of SlTCP14-2 localisation in non-infected cells and tissues challenged with *P. capsici* culture filtrate (CF) contrasted these results, as relocalisation was not observed. Collectively, these results suggest that SLTCP14-2 relocalisation is not attributable to (PAMP-triggered) defence responses, but rather *P. capsici* infection and effector activity.

**Figure 5.**
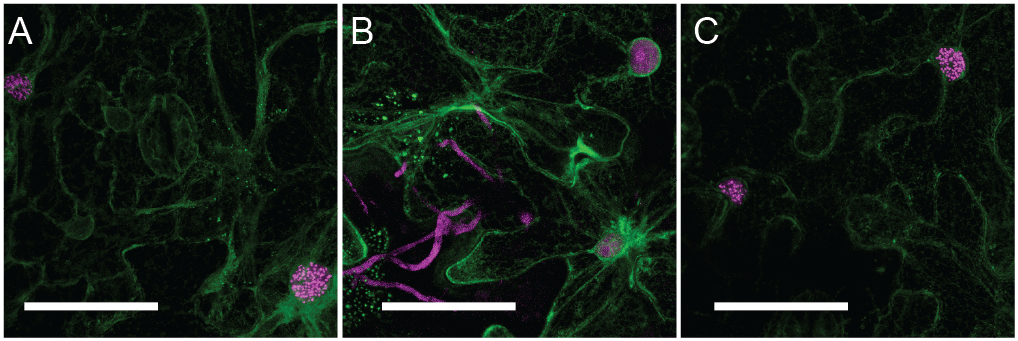
Speckled TCP14-2 (magenta) localisation can be observed in non-infected regions of the leaf (A), however the speckles disappear in tdTomato expressing *P. capsici* (magenta) infected regions (B). Infiltration with culture filtrate (C) does not have this effect. Green ER marker shows cell shape and viability. Scale bar = 50 um

### CRN12_997 dissociates TCP14-2 from chromatin

To test whether CRN12_997 interferes with SlTCP14-2 DNA binding properties we over-expressed CRN-SlTCP14 combinations and used a fractionation approach to separate chromatin-associated complexes from soluble proteins (Figure 6). Chromatin fractionation on cells only expressing SlTCP14-2 led to the co-purification of SlTCP14-2. Intriguingly, when co-expressed with CRN12_997, we observed a marked reduction in chromatin associated SlTCP14-2 (Figure 6). Consistent with the observed specificity between CRN12_997 and SlTCP14-2, CRN125_11 expression did not alter SlTCP14-2 levels in the chromatin associated fraction, strongly suggesting that CRN12_997 activity towards SlTCP14-2 causes its dissociation from chromatin.

**Figure 6.**
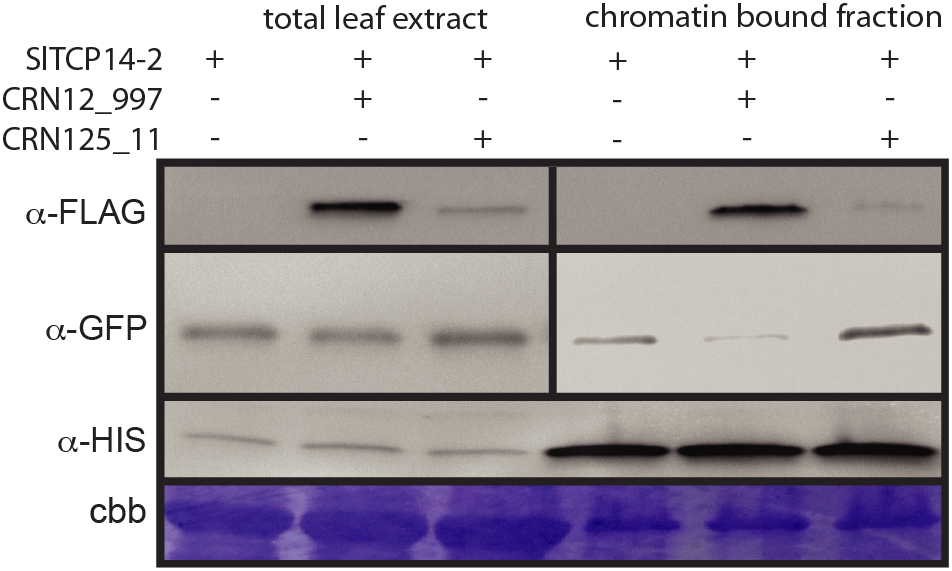
SlTCP14-2 signal is lower when co-expressed with CRN12_997. GFP-tagged SlTCP14-2 is expressed alone or co-expressed with FLAG-tagged CRN12_997 or CRN125_11. Top panel shows expression of both CRN proteins. An anti-Histone H3 blot and coomassie staining show chromatin enrichment and equal loading.

## Discussion

The plant immune signalling network comprises a vast and complex web of interconnected signalling cascades that collectively regulate defences against most microbes. Pathogens however compromise host immunity by interfering with key signalling components or “hubs” that regulate defence and enhance pathogen virulence. In this context, it may not be surprising that divergent pathogens such as nematodes, fungi and bacteria have independently evolved effectors that bind a common host target (6, 35, 36) or indeed, some effectors are conserved between pathogen species (36, 37). A remaining question is whether after speciation and a protracted co-evolutionary arms race, effector-interactions are preserved in divergent host-microbe systems.

Here we provide significant evidence suggesting that effector-target interactions identified between three *Hyaloperonospora arabidopsidis* CRN effector domains and the Arabidopsis TCP14 transcription factor, is preserved in the *P. capsici*-tomato interaction. The CRN effector protein family is diverse, but many CRN effector domains are conserved in divergent plant pathogenic oomycetes. We used this information to interrogate the literature for candidate CRN-protein interactions, and identified CRN-AtTCP14 interactions that were unveiled in a systems level study of effector-host target interactions in *Arabidopsis* (6). We expand on this initial work and show that CRN-host target interactions are conserved in the oomycete pathogen *P. capsici* and the Solanaceous host tomato. Expression of SlTCP14-2 in *N. benthamiana* leaves reduced *P. capsici* growth rates and onset of sporulation, suggesting that consistent with the view of TCP14 as important player in defence, this protein also appear to (positive) regulate immunity in tomato. Critically, co-expression of CRN12_997 and SlTCP14-2 restores *P. capsici* growth and sporulation.

In addition to our co-expression analyses, we present independent lines of evidence that suggest an interaction between CRN12_997 and SlTCP14-2. BiFC analyses demonstrated significant differences between CRN-SlTCP14 and control combinations, suggesting effector-target specificity for CRN12_997 and SlTCP14-2. We independently confirmed these observations as we could only simultaneously detect CRN12_997 and SlTCP14-2 in our Co-IP experiments.

To rationalise the interaction between CRN12_997 and SlTCP14-2 and its impact on *P. capsici* virulence and host immunity respectively, we investigated the activity of CRN12_997 towards SlTCP14-2 further. Co-localisation experiments revealed that in the presence of CRN12_997, tagRFP-SlTCP14-2 displayed a diffuse sub-nuclear re-localisation phenotype that contrasted normal speckled distribution patterns seen in cells expressing CRN125_11 and other control treatments (Figure 4). Given that nuclear speckles are generally assumed to be particles that have a role in transcription in both mammals and plants (37, 38), we hypothesised that re-localisation reflects interference of SlTCP14-2 function either caused by or resulting in dissociation of the target from nuclear chromatin. Indeed, chromatin fractionation assays showed that in the presence of CRN12_997, SlTCP14-2 levels drop in chromatin-bound fractions, suggesting a reduction in DNA binding affinity of SlTCP14-2 containing complexes. If true, these results would suggest that CRN12_997 binds an important transcriptional regulator of immunity in plants upon which activity is perturbed.

To our knowledge, this work represents the first example of a virulence strategy in filamentous pathogens that relies on the modification of host target DNA binding properties. Recent years has seen the emergence of evidence that suggests that effector mediated transcriptional reprogramming is crucial for pathogen success. TAL effectors from *Xanthomonas* can bind DNA directly in a sequence specific manner, activating the expression of host genes required for virulence in field conditions (39). With an extensive TAL effector family identified in bacteria, the presence of equivalent effector families in eukaryotes is yet to be demonstrated. Whether CRN12_997 requires an ability to bind DNA, its target or both, for virulence function is yet unknown. We routinely identified CRN12_997 in chromatin bound fractions in our assays, rendering both direct interactions as well as indirect DNA associations likely scenarios. In either case, alteration of SlTCP14-2 activity is likely to affect transcriptional changes, some of which may have been observed in tomato upon infection (34). Such events could not only underpin suppression of immune responses, but also participate in wholesale transcriptional reprogramming of host gene expression, linked to pathogen lifestyles (34). Given the importance of TCP14 in both Arabidopsis and solanaceous immune signalling, evidenced by the preservation of an effector-target interaction in divergent species, modification of TCP14 and its downstream targets in plants may allow an engineering strategy that result in crops that are more resistant to a wider range of pathogens.

## Acknowledgments

We would like to thank Dr. Andrew Howden and members of the Dundee Effector Consortium for constructive feedback on our work. This research was funded, in part, by the Biotechnology and Biological Sciences Research Council, the European Research Council and the Royal Society of Edinburgh.

## METHODS

### Homologies and alignments

HpaRXCRN sequences were obtained from Mukthar et al. (6). Tomato homologs for AtTCP14 and *P capsici* CRN homologs were identified using BLASTn, selecting Reciprocal Best Blast Hits (27, 40). The Hpa effectors were analysed using the methods described previously (28) to confirm that domain structure of both Hpa and Pc effectors was identical. Subsequently all other CRNs with similar domain structure were extracted from Haas et al (25). Sequences were truncated after the DWL domain and aligned using MUSCLE (41). A maximum likelihood tree was created with PhyML and visualised with TreeDyn, using the phylogeny.lirmm.fr webserver. A conservation motif was drawn using jalview (42). TCPs were aligned using MUSCLE. Tomato TCP expression data was obtained from microarray data generated from a *Phytophthora capsici*-tomato time course infection experiment (28, 34).

### Transient expression and localisation

For co-localisation GFP-expressing pB7WGF2 plasmids containing CRN C-terminal constructs and RFP-expressing pGWB461 plasmids containing TCP constructs were transformed into *Agrobacterium tumefaciens* strain AGL1. Transformants were grown on LB medium containing Rifampicin and Spectinomycin. A single colony was grown overnight and resuspended in infiltration buffer (10 mM MgCl, 150 µM Acetosyringone) to an OD600 of 0.1 for confocal microscopy or 0.5 other assays. The buffer was mixed 1:1 with buffer containing *Agrobacterium* carrying the silencing suppressor P19 and infiltrated in four to five week old *N. benthamiana* leaves. For BiFC CRN and TCP constructs were made using pCL112 and pCL113 vectors (43) and transiently expressed by AGL1 infiltration at an OD600 of 0.01. Plants were grown in a glasshouse under 16 hours light and temperatures of 26 °C by day and 22 °C by night. *N. benthamiana* leaves transiently expressing the construct of interest were harvested for microscopy 2 days after infiltration. To maintain cell viability and improve optical properties after detachment, the leaves were infiltrated with water before mounting on a microscope slide. Leaf samples were imaged using a Leica SP2 confocal microscope. Peak excitation/emission wavelengths were 488/509 nm for GFP, 561/607 nm for tagRFP and 514/527 nm for YFP.

### Protein extraction, Co-IP and western blot

Plant tissues were harvested at 3 dpi. Protein extractions were done using GTEN buffer (10% Glycerol, 25 mM Tris, 1 mM EDTA, 150 mM NaCl) supplemented with 2% PVPP, 10 mM DTT and 1X Complete protease inhibitor cocktail (Thermo Scientific). Samples were pulled down using antiFLAG M2 agarose (Sigma) or strepII-tag magnetic beads (44) (IBA GmbH) and run on 4-20% TGX PAGE gels (Biorad) before transfer to PVDF membranes using a Biorad Transblot system. Blots were blocked for 30 minutes with 5% milk in TBS-T (0.1% tween), probed with mouse anti-FLAG antibodies (Santa Cruz Biotech)(1:4000), StrepII-HRP antibody (Genscript)(1:5000) or mouse anti GFP antibody (Santa Cruz) (1:5000) in 5% milk in TBS-T. FLAG and GFP blots were secondarily probed with anti Mouse-HRP antibodies (1:20.000). All blots were washed 3-4 times in TBS-T for 5 minutes before incubation with Millipore Luminata Forte substrate. Blots were imaged on a Syngene G:BOX XT4 Imager.

### Chromatin fractionation

Tagged candidate proteins were overexpressed in *N. benthamiana* as described above. Leaves were harvested and ground in liquid nitrogen 3 days post infiltration. Ground leaf tissue was suspended in 10 ml ice cold buffer (10 mM PIPES pH6.8, 10 mM KCl, 1.5 mM MgCl_2_, 340 mM Sucrose, 10% glycerol, 0.5% Triton X-100, (1 mM DTT) and 1X SIGMAFAST protease inhibitor cocktail (Sigma)) filtered through Miracloth (Calbiochem). A 6 ml aliquot of this whole cell extract was taken and centrifuged at 3000 g for 10 minutes at 4 °C. 1 ml of TCA-a (10 ml acetone, 2 ml TCA (20% w/v in H_2_O), 8 µl β-mercaptoethanol) was used to resuspend the pellet, containing chromatin bound proteins. After brief vortexing, these samples were stored at −20 °C for 1 hour. Samples were centrifuged at 16000 g for 30 minutes at 4 °C and the resulting pellets were washed 3 times with a/β-me (10 ml acetone 8 µl β-mercaptoethanol) before resuspending in SDS loading buffer.

### Disease assays

Leaves were harvested 3 days after infiltration and collected in 1.5 cm deep collection trays containing wet tissue paper. *P. capsici* inoculation was done using 10 µL droplets of zoospore solution (500,000 spores per mL) of strain LT1534. The trays were stored in an incubator under 16 hours light and set at 26 °C by day and 22 °C by night. Lesion diameters were measured 2 days post inoculation as previously described (28, 45). Detached leaves from *N. benthamiana* plants expressing ER-targeted GFP were infected with droplets of *P. capsici* expressing tdTomato and imaged 24 hours after infection. CF treatment was imaged 4 hours post infiltration and prepared as described before (31).

